# A novel workflow to improve multi-locus genotyping of wildlife species: an experimental set-up with a known model system

**DOI:** 10.1101/638288

**Authors:** Mark A.F. Gillingham, B. Karina Montero, Kerstin Wihelm, Kara Grudzus, Simone Sommer, Pablo S.C. Santos

## Abstract

Genotyping novel complex multigene systems is particularly challenging in non-model organisms. Target primers frequently amplify simultaneously multiple loci leading to high PCR and sequencing artefacts such as chimeras and allele amplification bias. Most next-generation sequencing genotyping pipelines have been validated in non-model systems whereby the real genotype is unknown and the generation of artefacts may be highly repeatable. Further hindering accurate genotyping, the relationship between artefacts and copy number variation (CNV) within a PCR remains poorly described. Here we investigate the latter by experimentally combining multiple known major histocompatibility complex (MHC) haplotypes of a model organism (chicken, Gallus gallus, 43 artificial genotypes with 2-13 alleles per amplicon). In addition to well defined “optimal” primers, we simulated a non-model species situation by designing “naive” primers, with sequence data from closely related Galliform species. We applied a novel open-source genotyping pipeline (ACACIA) to the data, and compared its performance with another, previously published, pipeline. ACACIA yielded very high allele calling accuracy (>98%). Non-chimeric artefacts increased linearly with increasing CNV but chimeric artefacts leveled when amplifying more than 4-6 alleles. As expected, we found heterogeneous amplification efficiency of allelic variants when co-amplifying multiple loci. Using our validated ACACIA pipeline and the example data of this study, we discuss in detail the pitfalls researchers should avoid in order to reliably genotype complex multigene systems. ACACIA and the datasets used in this study are publicly available at GitLab and FigShare (https://gitlab.com/psc_santos/ACACIA and https://figshare.com/projects/ACACIA/66485).

## Introduction

A key challenge for molecular ecologists is that they frequently work on systems with limited to no knowledge of their genomes. This means that the development of a genotyping approach often relies on information from closely related species available in genetic databases. Furthermore, assessing and validating genotyping methods can be particularly challenging when the structure of the target region is unknown.

Multigene complexes, such as resistance genes (R-genes) and self-incompatibility genes (SI-genes) in plants, immunoglobulin superfamily and major histocompatibility genes (MHC) in vertebrates, and homeobox genes in animals, plants and fungi, among many others, are particularly challenging to genotype in non-model organisms. As a result of high sequence similarity from recent gene duplication events, polymerase chain reaction (PCR) primers will frequently bind across multiple loci leading to the amplification of multiple allelic variants [3–5, 30, 31, 49, 53]. Unspecific locus amplification may lead to several biases during PCR since 1) chimeric sequences (hereafter “chimeras”; which may arise because of incomplete extension of sequences during a PCR cycle which are subsequently completed with a different allele template) are likely to become more frequent as more loci are amplified within an amplicon simply because there will be more gene variants from which chimeras can be generated [29]; 2) amplification bias of some gene variants relative to others may occur because primers preferentially bind to some alleles/loci (hereafter referred to as “PCR competition”) [33, 53]. Creative solutions in primer design and in PCR conditions, such as using pooled primers instead of degenerate primers [33], reducing the number of cycles and modifying elongation steps of PCRs [21, 29, 52], can significantly reduce amplification bias. However, even after the application of such methods, PCR biases will nonetheless persist and may lead to genotyping errors because: 1) chimerias may be difficult to distinguish from valid recombinant gene variants (frequent in multigene complexes [10]), resulting in either PCR artefacts being falsely validated as a true allelic variants (type I errors, hereafter referred to as “false positives”) or in true allelic variants being falsely rejected as an artefact (type II errors, hereafter referred to as “allele dropout”) and 2) poorly amplified allelic variants may not be sequenced resulting in allele dropout, particularly when the number of sequences per amplicon (a set of sequences of a target region generated within a PCR) is low [4, 14, 30, 31, 53].

The recent rapid dissemination of next generation DNA sequencing (NGS) platforms has provided molecular ecologists with an exciting opportunity to tackle the parallelised genotyping of multiple markers in numerous species, since it has allowed the generation of thousands of sequences (termed “reads”) per amplicon, at a fraction of cost and time needed previously [3, 31, 53]. However, NGS platforms have their own limitations, the most relevant being the relatively high amount of sequencing errors generated in a typical sequencing run [16, 19, 34, 47, 53]. For instance, Illumina, currently the mainstream technology for NGS amplicon sequencing, report an error rate (primarily substitutions of base pairs) of ≤ 0.1% per base for ≥ 75-85% of bases (see Glenn [16] for details), although final error rates are likely to be much higher and can reach up to 6% [34]. Indeed, previous genotyping studies in multi-locus-systems (>10) reported average amplification and sequencing artefact rates of 1.5% to 2.5% per amplicon [43, 45, 50]. Therefore, PCR competition when amplifying multiple loci per amplicon means that sequences from some genuine allelic variants occur at a similar frequency to PCR artefacts or sequencing errors [4, 14, 30, 53]. In this scenario, poorly amplified alleles cannot be easily distinguished from artefacts during allele validation, leading to further false positives and allele dropout during genotyping.

The need to distinguish PCR and sequencing artefacts from valid allelic variants has led to the development of multiple bioinformatic workflows (i.e. a set of bioinformatic steps during processing of sequencing data which eventually leads to genotyping, hereafter referred to as a “genotyping pipeline”). While all genotyping pipelines rely to some degree on the assumption that artefacts are less frequent than genuine allelic variants, they vary in the approach used to discriminate poorly amplified allelic variants from artefacts. Genotyping pipelines for complex gene families have been extensively reviewed in Biedrzycka *et al* [4]. Recently developed pipelines cluster artefacts to their putative parental sequences thereby increasing the read depths of true variants [31, 42, 49, 55]. Currently, the most commonly used pipeline for MHC studies is the AmpliSAS web server pipeline [49]. After chimera removal, AmpliSAS uses a clustering algorithm to discriminate between artefacts and allelic variants, which take into account the error rate of a particular NGS technology and the expected lengths of the amplified sequences. This is achieved in a stepwise manner, whereby it first clusters the most common variant (according to specified error rates) and then moves on to the next most common variant, until no variant remains to be clustered. Microbiome studies, which typically amplify hypervariable regions of the 16S rRNA gene from very diverse bacterial communities within a single amplicon, have used a similar strategy to AmpliSAS, whereby potential artefactual variants are clustered to suspected parental sequences using Shannon entropy (referred to as “Oligotyping” [13]) or other similar clustering methods [2, 9].

Most of the amplicon genotyping pipelines for multigene families available to molecular ecologists have only been tested on non-model organisms for which the real genotype is unknown (but see Sebastian *et al* [49]). As a consequence, studies have frequently depended on repeatability of duplicated samples to justify genotyping pipeline reliability [4, 14, 31, 45, 49, 53]. However for a given set of PCR primers and sequencing technology, PCR and sequencing bias, and thus in turn the rate of false positives and allele dropout, will be consistently repeatable [4]. For instance, the high rate of Illumina substitution errors are known to be not random (see references within Sebastian *et al* [49]) and therefore variants which result from substitution errors are highly repeatable between amplicons [4]. Furthermore, while the generation of PCR and sequencing artefacts is well known, the precise relationship between artefacts and the number of alleles amplified within an amplicon for a given set of primers and sequencing technology has never been described. Yet, having a clear indication of this relationship is an important step in predicting what are the optimal pipelines settings (e.g. predicting error rates) for a given number of loci amplified within an amplicon. The latter can only be achieved by experimentally manipulating CNV of a priori known genotypes before PCR amplification and NGS sequencing.

In this study, we manipulated known combinations of the MHC alleles of a model organism (the chicken, Gallus gallus) as an example of a target multigene region of interest to molecular ecologists, in order to accurately quantify the effects of PCR and sequencing artefacts on genotyping pipelines. While we focus on the MHC hereafter, all methods and results are applicable to any multigene family. Like many multigene complexes, MHC genes are subject to multiple gene conversion, duplication and deletion [39–41] and MHC gene copies vary considerably across and even within a species (reviewed in [26]). Therefore, the number of MHC loci present in a non-model study system often remains unknown. For instance, MHC class IIB CNV was found to be as high as 21 in some passerine species, resulting in up to 42 allelic variants amplified within an amplicon and strong CNV between individuals [4]. In contrast, the chicken MHC B complex is unusually simple, leading it to be coined as a “minimal essential” system, with only two MHC class I loci and two MHC class II loci [23–25]. The latter is therefore an ideal system to validate MHC genotyping pipelines for the following reasons: 1.) the structure of the B complex is well known with well-defined primers in conserved regions; 2.) the well characterised B complex haplotype lineages can be used so that the expected MHC genotyping results are known prior to sequencing and genotyping and 3.) CNV within an amplicon can be experimentally engineered by combining DNA samples from multiple MHC B complex haplotypes.

In order to perform the genotyping of known chicken MHC haplotypes and extract data concerning PCR and sequencing artefacts at each step of the genotyping workflow, we developed and calibrated our own genotyping pipeline (named ACACIA for **A**llele **CA**lling pro**C**edure for **I**llumina **A**mplicon sequencing data). ACACIA is written in Python and it takes advantage of several previously published software dedicated to genomics (detailed in the methods), as well as the widely used Biopython library [11] to handle genomic data. We experimentally generated a MHC dataset with a range of CNVs by combining DNA samples from multiple chicken MHC B complex haplotypes. Since MHC B complex in chickens is well characterised, optimal primers to amplify the entire exons which code for the antigen binding regions have been developed within the introns [17, 51]. However in most wildlife species, such extensive genomic information around the region of interest is unavailable. In order to avoid the problems associated with overfitting ACACIA to one specific dataset and also in order to replicate the challenge of designing primers for a non-model species, we additionally designed primers within the exons coding for antigen-binding regions using sequence data from closely related Galliform species that were not chickens (hereafter referred to as “naive primers”). The latter enabled us to gain insight into the relative amount of artefacts generated by an intentionally sub-optimal set of primers, for which we expected allele dropout.

Specifically, this study aimed to:

1. validate ACACIA using experimentally manipulated genotypes with different CNV that are known a priori;
2. accurately describe the relationship between PCR/sequencing artefacts and CNV by experimentally varying CNV and primer design in a model system.

## Materials and Methods

### Samples and DNA extraction

Chicken blood samples originated from experimental inbred lines kept at the Institute for Animal Health at Compton UK (lines 72, C, WL and N) and the Basel Institute for Immunology in Basel Switzerland (lines H.B15 and H.B19+), as detailed in Jacob *et al* [20], Shaw *et al* [51] and Wallny *et al* [58]. These lines carry seven common B haplotypes: B2 (line 72), B4 and B12 (line C), B14 (line WL, sometimes referred as W), B15 (H.B15), B19 (H.B19) and B21 (line N). All the lines are homozygotes at the MHC except line C, which was not used in this study. In each haplotype are two class II B loci: BLB1 (previously known as BLBI or BLBminor) and BLB2 (BLBII or BLBmajor), with alleles now designated as BLB1*02 and BLB2*02 from the B2 haplotype, etc. All alleles have different nucleotide sequences, except BLB1*12 and BLB1*19. DNA was isolated from blood cells by a salting out procedure [37].

### Generating 41 artificial MHC genotypes

We artificially generated 43 genotypes of varying CNV by combining equimolar amounts of DNA samples from the seven MHC haplotypes mentioned above (Table 1; created genotypes listed in Supplementary Table S1).

### Optimal primers for chicken MHC Class II

We targeted the entire 241 bp of exon 2 of MHC class II, the polymorphic region known to code for antigen binding sites, using the primers OL284BL (5’-GTGCCCGCAGCGTTCTTC-3’) and RV280BL (5’-TCCTCTGCACCGTGAAGG-3’) [17]. The primers are not locus specific and bind to both loci of the chicken B complex.

### Naive primer design for chicken MHC Class II

In order to naively design primers, we downloaded 61 exon 2 MHC Class II sequences from seven Galliform species (*Coturnix japonica*, *Crossoptilon crossoptilon*, *Meleagris gallopavo*, *Numida meleagris*, *Pavo cristatus*, *Perdix perdix* and *Phasianus colchicus*) from the GenBank (https://www.ncbi.nlm.nih.gov/genbank/). We then used Primer3 [48, 57] to design the forward primer GagaF1 (5’-WTCTACAACCGGCAGCAGT-3’) and the reverse primer GagaR2 (5’-TCCTCTGCACCGTGAWGGAC-3’) aiming at amplifying 151 bp of exon 2.

**Table 1.** The number of alleles per genotype, the number of genotylongwith a certain number alleles and the number of amplicons with a certain number alleles (all genotypes were duplicated) for the chicken datasets used in this study. The list haplotypes used to artificially create the genotypes are listed in Supplementary Table S1.

| Number of alleles per genotype | Number of genotypes | Number of amplicons |
| --- | --- | --- |
| 2 | 7 | 14 |
| 4 | 7 | 14 |
| 6 | 7 | 14 |
| 8 | 7 | 14 |
| 10 | 7 | 14 |
| 11 | 5 | 10 |
| 12 | 2 | 4 |
| 13 | 1 | 2 |
| **Total** | **43** | **86** |

### PCR Amplification, Library Preparation, and High-Throughput Sequencing

For all datasets we replicated all individuals in order to estimate repeatability (n_individuals_ = 43 and n_amplicons_ = 86). Individual PCR reactions were tagged with a 10-base pair identifier, using a standardised Fluidigm protocol (Access Array™ System for Illumina Sequencing Systems, ©Fluidigm Corporation). We first performed a target specific PCR with the CS1 adapter and the CS2 adapter appended. To enrich base pair diversity of our libraries during sequencing, we added four random bases to our forward primer. The CS1 and CS2 adapters were then used in a second PCR to add a 10bp barcode sequence and the adapter sequences used by the Illumina instrument during sequencing.

The first PCR consisted of 3–5 ng of extracted DNA, 0.5 units FastStart Taq DNA Polymerase (Roche Applied Science, Mannheim, Germany), 1× PCR buffer, 4.5 mM MgCl_2_, 250 μM of each dNTP, 0.5 μM primers, and 5% dimethylsulfoxide (DMSO). The PCR was carried out with an initial denaturation step at 95°C for 4 min followed by 30 cycles at 95°C for 30 s, 60°C for 30 s, 72°C for 45 s, and a final extension step at 72°C for 10 min. The second PCR contained 2 μl of the product generated by the initial PCR, 80 nM per barcode primer, 0.5 units FastStart Taq DNA Polymerase, 1× PCR buffer, 4.5 mM MgCl_2_, 250 μM of each dNTP, and 5% dimethylsulfoxide (DMSO) in a final volume of 20 μl. Cycling conditions were the same as those outlined above but the number of cycles was reduced to ten.

PCR products were purified using an Agilent AMPure XP (Beckman Coulter) bead cleanup kit. The fragment size and DNA concentration of the cleaned PCR products were estimated with the QIAxcel Advanced System (Qiagen) and by UV/VIS spectroscopy on an Xpose instrument (Trinean, Gentbrugge, Belgium). Samples were then pooled to equimolar amounts of DNA. The library was prepared as recommended by Illumina (Miseq System Denature and Dilute Libraries Guide 15039740 v05) and was loaded at 7.5 pM on a MiSeq flow cell with a 10% PhiX spike. Paired-end sequencing was performed over 2 × 251 cycles.

### Data analysis with the ACACIA pipeline

ACACIA consists of 11 consecutive steps of data processing.. The software requires two non-standard python libraries (Pandas [35] and Biopython [11]) as well as six third-party software (FastQC (www.bioinformatics.babraham.ac.uk/projects/fastqc/), FLASh [32], VSEARCH [46], BLAST [1], MAFFT [22] and Oligotyping [13]), which can all be installed with one command. The input files are any number of FASTq files, which are the current canonical output of the Illumina platform. The step-by-step workflow is described below:

1. **Generating Quality Reports.** Sequencing quality is assessed for each FASTq file yielded by the sequencing platform, with the FastQC tool. Reports for each file are produced in HTML format for visual inspection.
2. **Trimming low quality ends of forward and reverse reads (optional).** The information generated in step #1 is crucial for an informed decision about how many (if any) bases should be trimmed out of each read. If trimming is performed here, step #1 is repeated. Shorter FASTq files are generated as output of this step.
3. **Merging paired-end reads (optional).** This concerns projects with paired-end sequencing only and should be skipped if using data from single-end sequencing (note: the names of the paired forward and reverse FASTq files should be identical prior to the first “_” character, e.g.: ID1S1L001_R1_001.fastq and ID1S1L001_R2_001.fastq). The reads of file pairs are merged using FLASh [32]. The minimum and maximum lengths of overlap during merging can be adjusted by the user to improve performance (defaults are zero and read length, respectively). New FASTq files with merged sequences are generated as output, as well as a series of .log files which allow users to monitor merging performance.
4. **Trimming primers.** After prompting users to enter the sequences of the primers used for target amplification, ACACIA trims primer sequences from both ends of the merged sequences (IUPAC nucleotide ambiguity codes are allowed). Primerless sequences are written into FASTq files which are the output of this step. The Python functions for trimming primers and low-quality ends (step #2) are part of the core ACACIA pipeline. External tools were avoided here to decrease dependency on further software.
5. **Quality-control.** Users are then prompted to enter the values of two parameters (q and p) in order to filter sequences based on their mean phred-scores. First, q stands for quality and denotes a phred-score threshold that can take values from 0 to 40. Second, p stands for percentage and denotes the proportion of bases, in any given sequence, that have to achieve at least the quality threshold q for that sequence to pass the quality filter. ACACIA uses the default values q = 30 and p = 90 if users do not explicitly change them. In practical terms, these thresholds correspond to an error probability lower than 10-3 in at least 90% of bases for each sequence. All information on quality data of sequences passing this filter is then removed and FASTA files with high-quality sequences are given as the output of this step.
6. **Removing singletons.** A large proportion of sequences contain random errors inherent to the sequencing technology [44]. In order to decrease file sizes without risking loss of relevant allele information, ACACIA removes all singletons (sequences that appear one single time) in an individual amplicon.
7. **Removing chimeras.** The chimera identification tool VSEARCH [46] is employed here, with slightly altered settings (alignwidth = 0 and mindiffs = 1) aiming at increasing sensitivity to chimeras that diverge very little from one of the “parent” sequences. FASTA files with non-chimeric sequences, along with log files for each individual amplicon, are given as output.
8. **Removing unrelated sequences.** All remaining sequences are then compared with a set of reference sequences chosen by users. This step aims at removing sequences that passed all filters so far but are products of unspecific priming during PCR. Typically, sequences phylogenetically related to those being analyzed can be downloaded from the GenBank (www.ncbi.nlm.nih.gov/genbank/). Users are prompted to provide one FASTA file with reference sequences, which is converted by ACACIA to a local BLAST database [1] and used for BLAST. Only sequences yielding high-scoring hits to the local database (expectation value threshold = 10) are written into new FASTA files as an output of this step, which is the workflow’s last filtering procedure.
9. **Aligning.** The MAFFT aligner [22] is used to perform global alignments of sequences that have passed filters. Since all sequences are pooled into one single alignment output file, the individual IDs are now transferred from file names into the FASTA sequence headers. We have successfully aligned up to 603,513 sequences in a desktop computer with four CPUs and 32GB of RAM. Users with a significantly higher number of sequences might find it useful to increase the computational parallelization of the aligner as described recently [38].
10. **Calling candidate alleles.** The Oligotyping tool [13] is used to call candidate alleles. Although originally conceived as a tool for identifying variants from microbiome 16S rRNA amplicon sequencing projects, we recognised Oligotyping as ideal for other forms of highly variable amplicon sequencing projects. This step consists of concatenating high-information nucleotide positions (defined by entropy analysis of the alignment produced in the previous step) and subsequently using entropy information to cluster divergent variants, while grouping redundant information and filtering out artefacts. Although Oligotyping was conceived as a supervised tool, we automated the selection of parameter values aiming at high tolerance. This has the advantage of running an unsupervised instance of Oligotype as a pipeline step, at the cost of keeping potential false positives among the results. Report files with a list of candidate alleles grouped by individual amplicons are the output of this step.
11. **Allele calling and final reporting.** A Python script is used to perform the final allele calling by filtering out Oligotyping results according to the following criteria:

- Removal of unique allele variants (Y/N). Setting Y (yes) removes all alleles identified in one single individual amplicon;
- Absolute number of reads (abs_nor): minimum number of sequences that need to support an allele, otherwise the allele is considered an artefact. Ranges between 0 and 1000, with default = 10;
- Lowest proportion of reads (low_por): in order to be called in an individual amplicon, an allele needs to be supported by at least the proportion of reads, within that individual amplicon, that is declared here. Ranges between 0 and 1, with default = 0, while a value greater than 0 is recommended for data sets with ultra deep sequencing depth, which can suffer more from false positives [4].

Subsequently, putative alleles with very low frequency (both at the individual and population level) are scrutinised again. If the proportion of reads of a putative allele within an individual amplicon is less than 10 times lower than the next higher ranking allele, and if it is very similar (one single different base) to another, more frequent allele present in the same individual amplicon, that putative allele is considered an artefact and removed. Finally, if an individual amplicon has fewer than 50 sequences following all of the allele calling validation steps, it is eliminated. Users are able to change all parameter values, but ACACIA recommends settings based on our benchmarking. The output of this step consists of four files:

- **allelereport.csv**: a brief allele report listing genotypes of all individual amplicons as well as frequencies and abundances of all alleles found in the run;
- **allelereport_XL.csv**: a detailed allele report including the number of reads supporting each allele both within individuals and in the population;
- **pipelinereport.csv**: a pipeline report quantifying read counts and sequences failing or passing each pipeline step described above;
- **alleles.fasta**: a FASTA sequence file of all alleles identified in the run.

We investigated the best abs_nor and low_por for our datasets by first looking at the allele calling accuracy (the proportion of alleles that have been correctly called) and repeatability (the proportion of alleles, including false positives, called in both PCR replicates) at varying abs_nor values (range: 0-40, with low_por set at 0) first, and at varying low_por values (range: 0-0.02, with the optimal abs_nor, in our case 10) second. The latter is how we recommend users to find their optimal settings, although the range of abs_nor and low_por values to be investigated may vary across different datasets, depending on where the “peak” optimal setting lies.

The pipeline is supervised by a configuration text file (config.ini) which is appended every time users enter one of the settings mentioned above. Users can avoid running ACACIA interactively (and run the whole workflow in a “hands-free” mode) by providing a complete config.ini file at the beginning of the workflow. A template of a config.ini file is given in ACACIA’s repository (https://gitlab.com/psc_santos/ACACIA/blob/master/config.ini).

### Data analysis with the AmpliSAS pipeline

To compare how ACACIA performed relative to an existing relevant pipeline, we applied the web server AmpliSAS pipeline to our chicken datasets [49]. The default AmpliSAS parameters of a substitution error rate of 1% and an indel error rate of 0.001% for Illumina data was used. We then tested for the optimal ‘minimum dominant frequency’ clustering threshold for a given filtering threshold (i.e. 0.5% for the ‘minimum amplicon frequency’), by testing a set of thresholds of 10%, 15%, 20% and 25%. All clustering parameters tested gave an allele calling accuracy of 97%, but we chose the 25% clustering threshold because it was the only parameter which resulted in no false positives.

Subsequently, AmpliSAS filters for clusters that are likely to be artefacts, including chimeras and other low frequency artefacts that have filtered through the clustering step [49]. The default setting for the filtering of low frequency variants (i.e. ‘minimum amplicon frequency’) is 3%. However this value was far too high for our datasets, and we tested a range of filtering threshold between 0% and 1% at 0.1% intervals (i.e. 0%, 0.1%, 0.2% etc.). We assessed the optimal filtering threshold using both allele calling accuracy and repeatability.

## Results

### Sequencing depth for each dataset and proportion of artefacts detected using ACACIA

A total of 530,101 paired-end reads were generated for the optimal primers dataset, which amounted to an average of 6,164 reads per amplicon (n = 86). For the naive primers dataset, 994,338 paired-end reads were generated, amounting to an average of 11,562 reads per amplicon (n = 86). The proportion of artefacts identified at each step of the ACACIA pipeline for the chicken datasets combined is illustrated in Figure 1. Workflow filtering removed the highest proportion of reads when filtering for singletons (13.6%) and chimeras (14.2%). After all filters, 66.4% of the original raw reads were used for allele calling.

**Figure 1.**
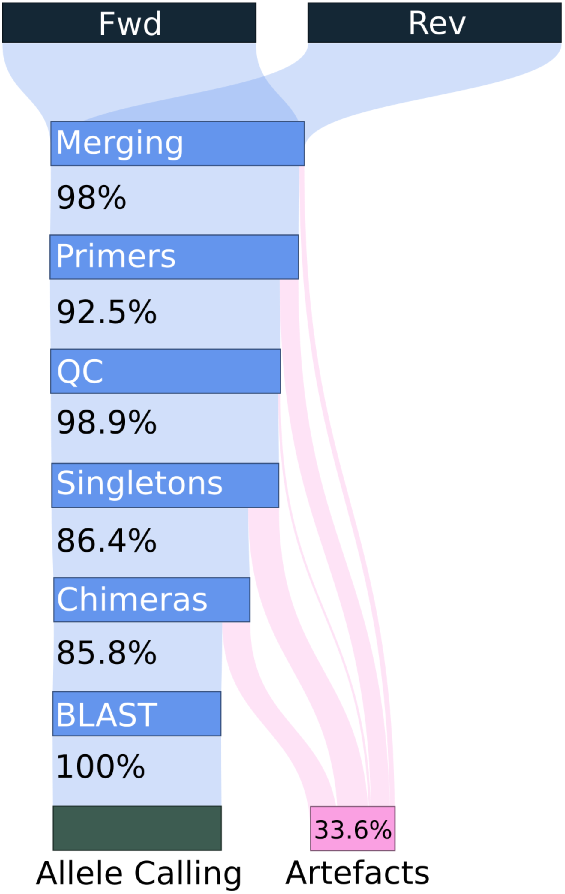
Flow diagram of reads and sequences from two Illumina runs analysed with ACACIA. Blue bars correspond to filters, and the percentages given correspond to the sequences kept at each step for further analyses. The percentage given at the bottom for artefacts refers to the total amount of reads in the beginning of the process. (Fwd & Rev) raw forward and reverse reads; (Mrg) paired-end read merger; (Prm) primer filter; (QC) quality control; (Sgt) Singleton removal; (Chm) chimera removal; (Blt) BLAST filter.

### Optimal settings of different workflows

We compared allele calling repeatability optimal abs_nor and low_por settings when using the ACACIA workflow. We first fixed the abs_nor setting at 10 and tested different low_por values and found that the optimal setting was 0 across both datasets (Figure 2a.). Lower low_por values increased allele dropout. We then tested the optimal abs_nor setting for a fixed low_por value of 0 and found that the optimal setting was 10 across both datasets (Figure 2b.). An abs_nor value of 0 increased the rate of false positives and whilst a value above 10 increased the rate of allele dropout.

For the AmpliSAS workflow, we investigated the optimal filtering threshold and found differing optimal values between datasets. For the optimal primer dataset we found that the optimal filtering threshold was 0.3 whilst 0.5 was found to be optimal for the naive dataset (Figure 2c.).

**Figure 2.**
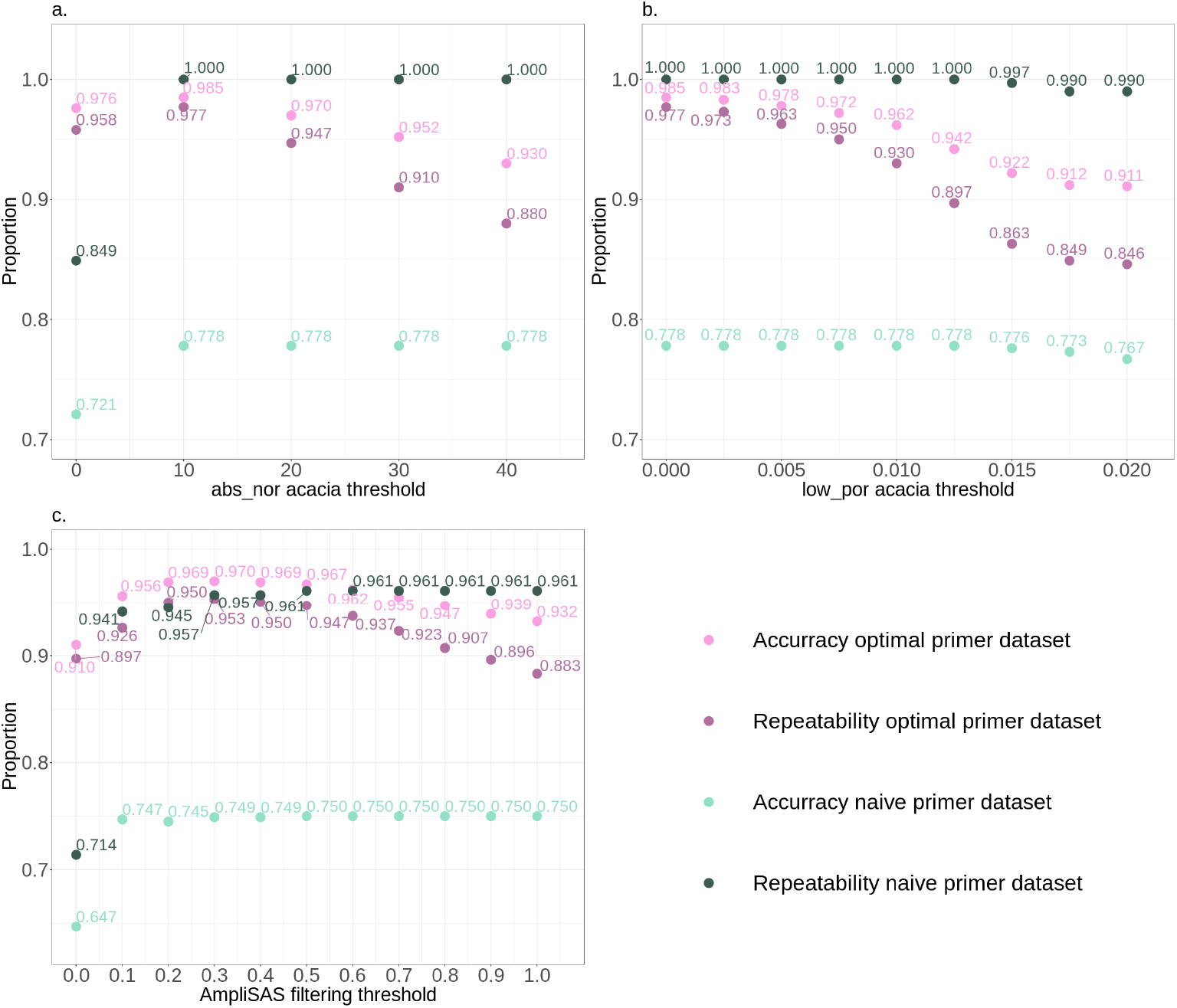
Allele calling accuracy and repeatability for the two datasets of this study (optimal primer or naive) at different abs_nor threshold settings with low_por set at 0 within the ACACIA pipeline (a.); at different low_por threshold settings with abs_nor set at 100 within the ACACIA pipeline (c.); and, at different filtering thresholds (i.e. ‘minimum amplicon frequency’) within the AmpliSAS pipeline.

### AmpliSAS vs ACACIA: optimal primers dataset

When using the optimal settings of the ACACIA workflow, comparison of results with expected genotypes revealed that nine alleles dropped out, no false positives were found (Table 2) and allele calling accuracy was 98.5% (Figure 2a. and b.). All instances of allele dropout derived from the B21 haplotype. For two genotypes, both BLB2*21 and BLB1*21 dropped out. For four genotypes, only BLB1*21 dropped out and for one genotype only BLB2*21 dropped out (Table 2). Allele calling repeatability was 97.7%.

Using the optimal settings in AmpliSAS, 17 alleles dropped out, one false positive was found (Table 2) and allele calling accuracy was 97% (Figure 2c.). As with ACACIA, most allele dropouts (16 of 17) derived from the B21 haplotype. For three genotypes, both BLB2*21 and BLB1*21 dropped out. For nine genotypes, only BLB2*21 alleles dropped out and for one genotype only BLB1*21 allele dropped out. Finally for one genotype the allele dropout was BLB2*04 and the same genotype had a false positive allele (Table 2). Allele calling repeatability was 95.3%.

**Table 2.**
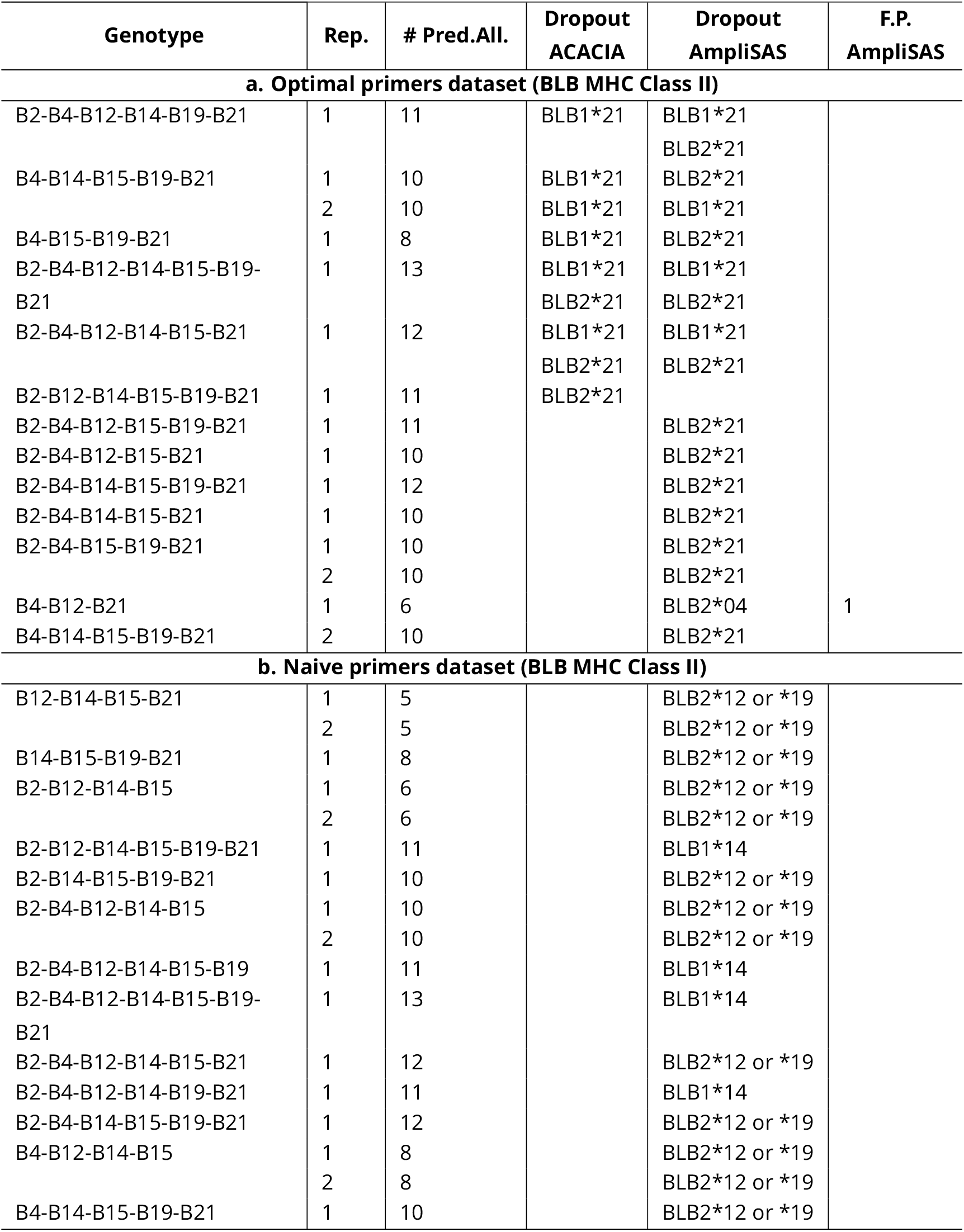
Genotypes with predicted number of alleles (# Pred.All.), allele dropouts (Dropout) and false positives (F.P.) using ACACIA and AmpliSAS (excluding allele dropout due to primer mismatch in the naive primers dataset).

### AmpliSAS vs ACACIA: chicken naive primers dataset

Using the optimal settings of ACACIA, we found 134 allele dropouts and allele calling accuracy was 77.8% (Figure 2a. and b.). However, all dropouts were from the alleles BLB2*04, BLB2*15 or BLB2*21, for which a primer mismatch was present. Therefore, all allele dropouts could be explained by primer design and allele calling repeatability between both replicates was 100%. Using the optimal settings of AmpliSAS, we found 152 allele dropouts and allele calling accuracy was 75.2% (Figure 2c.). As above, 134 dropouts were due to a mismatch with the forward primer. The remaining 17 alleles that dropped out were BLB2*12 or *19 (13 alleles) and BLB1*14 (4 alleles). Allele calling repeatability between both replicates was 96.1%.

### Relationship between number of alleles amplified and artefacts

The proportion of sequences classified as artefacts was much higher for PCRs using the optimal primer set than when using the naive primer set (Figure 3a. and 3b.). For all chicken data sets, when considering non-chimeric artefacts, there was a positive relationship between the proportion of artefacts and the number of alleles amplified (Figure 3a.). There is a logarithmic relationship between the proportion of chimeric artefacts and the number of alleles amplified whereby the proportion of chimeric reads no longer increased with number of alleles amplified when amplifying more than 4-6 alleles (Figure 3b.). The total number of unique chimeric reads also tended to follow a logarithmic relationship, whereby the number of unique chimeric variants seemed to no longer increase with the number of alleles amplified when amplifying more than 10 alleles (Figure 3c.). The total number of parental variants generating chimeras also did not increase with CNV when amplifying more than six alleles (Figure 3d.). Finally, the contribution of allelic variants to the proportion of reads decreased sharply with increasing number of alleles when amplifying less than 4-6 alleles (Figure 4). However the contribution of allele variants to the proportion of reads stabilised when amplifying more than 4-6 alleles (Figure 4). Both alleles from the B21 haplotype in the optimal dataset and the BLB1*04 allele in the naive dataset consistently amplified poorly when co-amplifying with alleles from other haplotypes (Figure 4).

**Figure 3.**
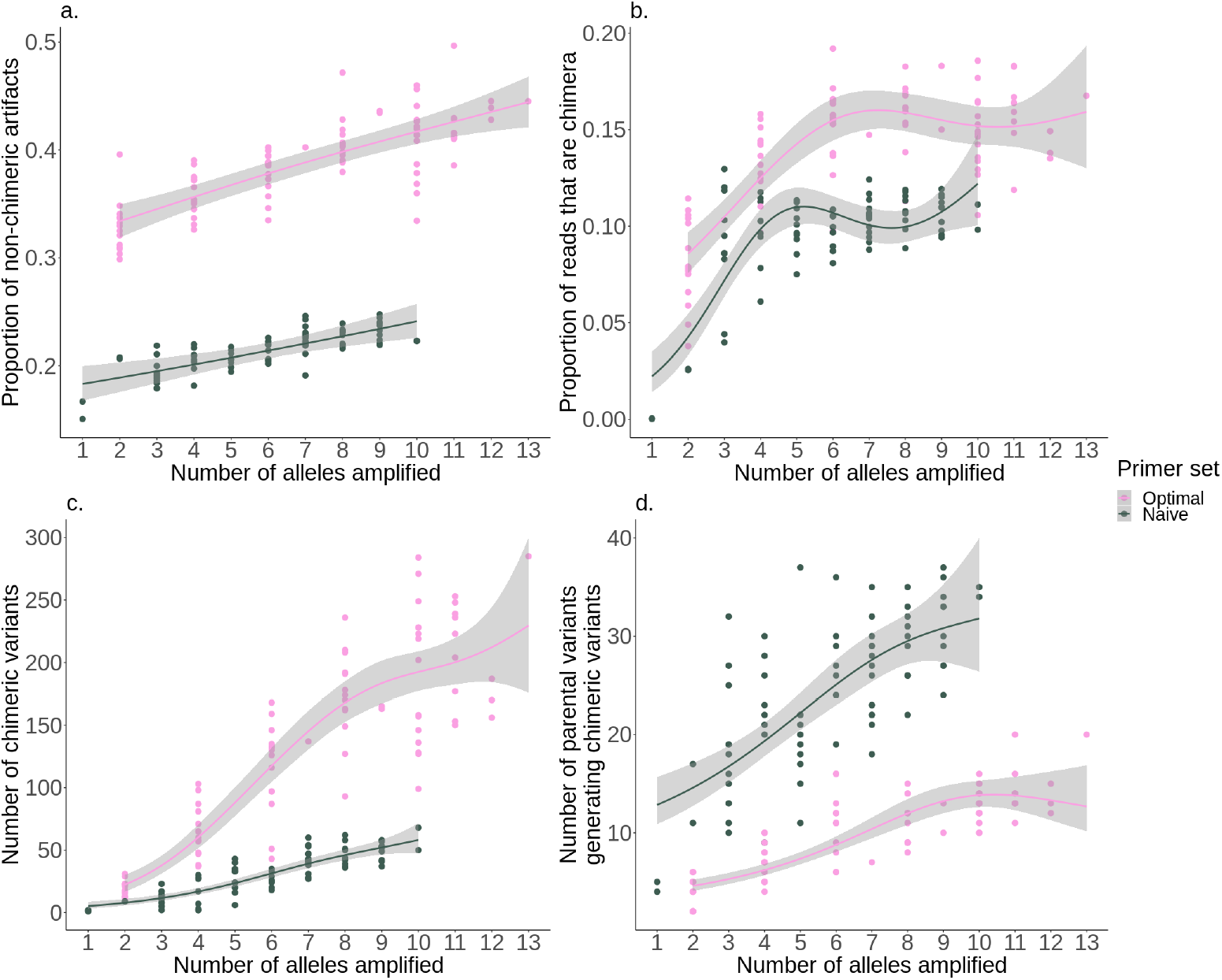
The relationship between the number of alleles amplified and: the proportion of non-chimeric reads (a.); the proportion of chimeric reads (b.); the absolute number of chimeric variants (c.); and, the absolute number of parental variants generating chimeric reads (d.). All relationships were fitted with general additive model using the ggplot package [59] in R [56] using a binomial distribution for (a.), (b.) and (f.), and a Poisson distribution corrected for over-dispersion for (c.) and (d.).

**Figure 4.**
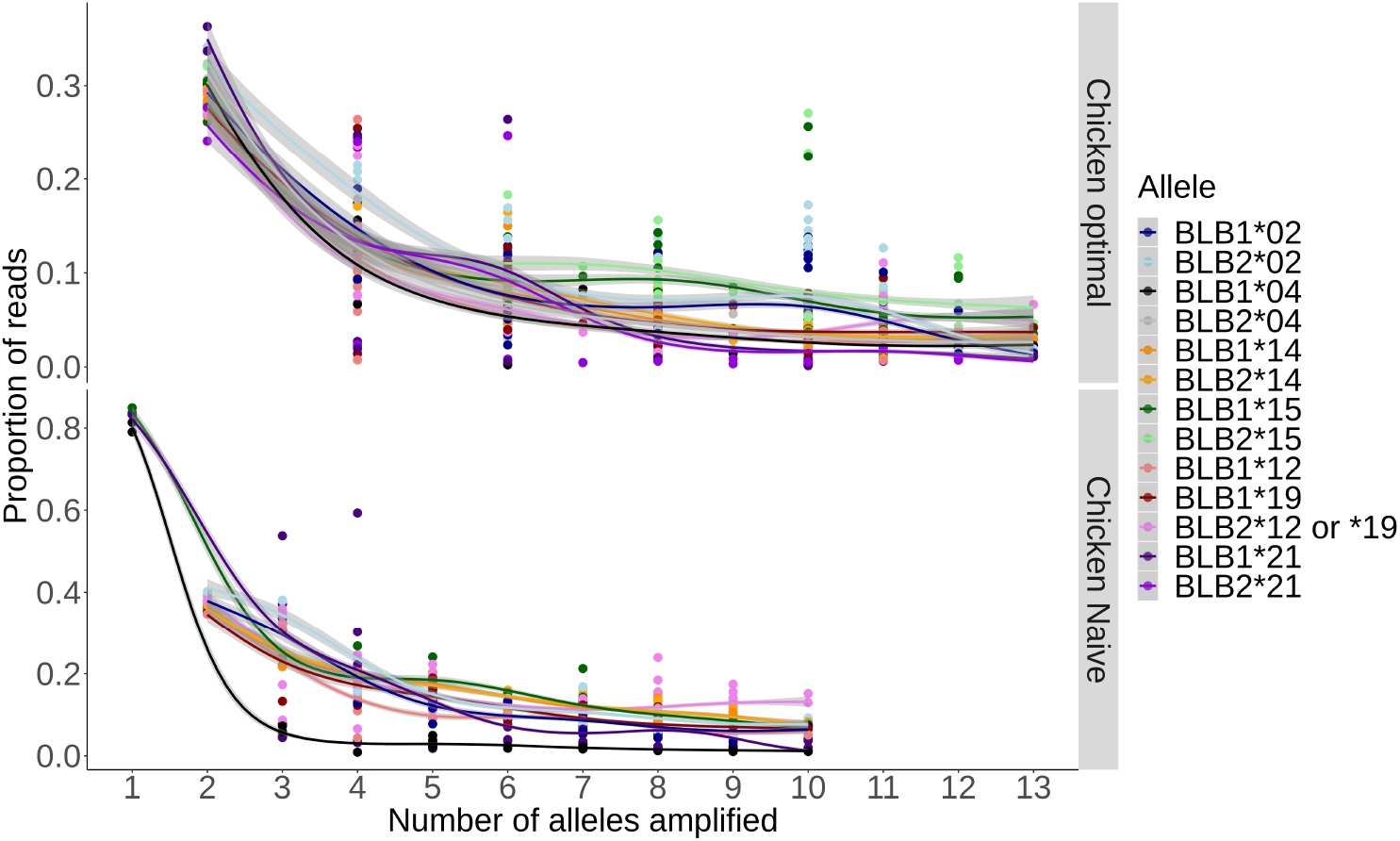
The relationship between the number of alleles amplified and the proportion of reads for each real allelic variant. All relationships were fitted with general additive model using the ggplot package [59] in R [56] using a binomial distribution.

## Discussion

Using known MHC genotypes for two datasets (chicken MHC Class II B complex), we achieved high allele calling accuracy (≥98.5%) and repeatability (≥97.7%) using ACACIA. With fewer allele dropouts and false positives, the ACACIA pipeline performed better than AmpliSAS. We demonstrated the “costs” of designing primers within MHC exon 2 in terms of allele dropout, with three common alleles failing to amplify when using primers naively designed from sequences of related Galliform species. We also explored the relationship between artefacts and CNV, and found that surprisingly, the relationship between the proportion of chimeric artefacts and CNV was not linear but rather leveled when amplifying more than 4-6 alleles. However, non-chimeric artefacts did increase linearly with increasing CNV. As expected we found heterogeneous amplification efficiency of allelic variants when amplifying multiple loci within a PCR. Below we discuss in further detail the ACACIA, AmpliSAS and other genotyping pipelines, primer design for non-model organisms, the relationship between CNV and artefacts, the effect of chimera formation on genotyping pipelines and, finally, we conclude by advising users on important points to consider when genotyping complex multigene systems in non-model organisms.

### AmpliSAS vs ACACIA

Experimentally generating CNV of known chicken MHC class II genotypes allowed us to validate our ACACIA pipeline to genotype systems with high CNV complexity at high accuracy and repeatability across replicates. While we achieved higher allele calling accuracy and repeatability using ACACIA than the AmpliSAS web server pipeline, we do not claim that ACACIA will necessarily perform better than AmpliSAS with all datasets. To demonstrate the latter we would need to test both pipelines on a larger number of datasets and/or on simulated datasets. In addition, while our pipeline should suit data generated with any next-generation sequencing technologies, we have only tested ACACIA with paired-end Illumina sequencing technology.

The most apparent benefit of using the AmpliSAS web server is that it is relatively easy to use for users with limited knowledge of scripting languages (such as PYTHON, PERL, C++ or R). However, we have noticed that a number of studies report using default settings when applying the AmpliSAS pipeline to their dataset. We find this concerning since, as our study demonstrates, the default clustering and filtering parameters are unlikely to be optimal for most datasets. Indeed, allele calling accuracy was much lower when using the default settings (81.8%) as compared to the optimal settings (97%) in the optimal primer dataset in our study, due to high allele dropout when using the default settings. We therefore strongly discourage users from using default settings and advise to permutate between different filtering and clustering parameters in order to find the best settings when using the AmpliSAS pipeline.

An important disadvantage of the AmpliSAS web server is that at the time of writing, sequencing depth per amplicon was limited to 5,000 reads. The latter is particularly problematic when wishing to genotype systems with complex CNV, which require high sequencing depth to genotype with high repeatability [4]. For datasets with sequencing depth above 5000 reads, AmpliSAS can be run locally but we found that, unlike the web server, the local version of AmpliSAS had limited documentation and troubleshooting was time consuming.

Once installed, ACACIA does not require users to have experience with scripting languages, allows genotyping with virtually unlimited sequencing depth and provides output data reporting the number of reads kept at each step of the pipeline. The latter should aid users when deciding upon optimal parameters and thresholds. As for the AmpliSAS pipeline, we advise to not use default parameters of ACACIA without critically assessing different parameters for each dataset. In particular, we urge users to permutate between different settings of abs_nor and low_por parameters. We advise to first search for the optimal abs_nor setting with a fixed low_por parameter of 0 because it is likely that it is only necessary to change the low_por parameter setting from 0 in datasets with ultra deep sequencing depth. If it is subsequently found that the optimal low_por setting is greater than 0, users should repeat the permuting step of abs_nor until the optimal settings are found. Of course finding optimal settings requires the inclusion of replicates for at least a subset of the dataset. We therefore recommend that a sufficient number of replicates are always included in genotyping runs to obtain sufficiently accurate repeatability values.

### Comparing ACACIA to other pipelines

Prior to the development of AmpliSAS and ACACIA, researchers who wished to genotype complex multigene systems generally relied on either earlier software such as SESAME [36] or jMHC [54] or their own customised scripts (e.g. [27, 61]). However while both SESAME and jMHC aided allele calling workflows by allowing users to demultiplex sequences and to generate tables which contains sequence variants and the number of reads, they do not allow users to apply an automated workflow to distinguish artefacts from real allelic variants.

Genotyping pipelines have evolved and matured in the last decade, however all genotyping pipelines rely to some degree on the assumption that artefacts are in general less frequent than genuine allelic variants. However genotyping pipelines vary in the methods used to discriminate poorly amplified allelic variants from artefacts. An early pipeline suggested by Radwan *et al* [45], which expanded from initial pipelines suggested by Kloch *et al* [27] and Zagalska-Neubauer *et al* [61], set a threshold below which all variants are considered artefacts (e.g. <1.5% per amplicon in Radwan *et al* [45]). This threshold is set by comparing rare variants to more common variants within an amplicon to determine whether the rare variant can be explained as an artefact (i.e. 1 to 2 bp mismatch compared to a common variant within an amplicon or a PCR chimera from two common parental variants within an amplicon). The weakness of this genotyping pipeline is that it relies on a single threshold below which all variants are considered artefacts, potentially making it particularly vulnerable to allele dropout [53]. A second method was suggested by Sommer, Courtiol, & Mazzoni [53], which relied on comparisons between duplicated amplicons and a series of decision making trees to discriminate between allelic variants and artefacts. While the pipeline of Sommer, Courtiol, & Mazzoni also assumes that artefacts are less frequent than most allelic variants, it does not rely on a single threshold below which all sequences are considered artefacts. However, one potential weakness of this method is that it may be more vulnerable to repeatable artefacts and thus to false positives, particularly in systems highly diverse in terms of high copy number variation (CNV>10 [4]).

A further disadvantage of all the above early genotyping pipelines is that much of the sequencing depth data is wasted by simply discarding low threshold sequences. In order to maximise the available sequencing depth, recent genotyping methods have clustered artefactual (non-chimeric) sequences to their suspected parental variant to increase genotyping confidence. This trend has been particularly strong in the 16S rRNA microbiome community, which have traditionally clustered sequence variants to so called operational taxonomic units (OTUs) using a fixed similarity threshold (usually 97% similarity). More recent 16S rRNA clustering methods such as the entropy based Oligotyping tool used within ACACIA [13], as well as model based methods such as DADA2 [9] and Deblur [2], have used alternative and more sophisticated statistical methods to simple similarity thresholds to distinguish sequence variants that differ by as little as one base pair. The clear benefit of clustering is that it significantly reduces the number of reads with low abundances, while increasing the read counts from poorly amplified allelic variants. However even the most sophisticated clustering methods will retain some artefacts within datasets [2, 8, 13], hence the need for additional filtering steps following clustering. Downstream filtering strategies can also resemble the pre-clustering pipelines strategies mentioned above as was applied by Biedrzycka *et al* [4] using AmpliSAS in a highly complex system (19 to 42 allelic variants per amplicon). Biedrzycka *et al* [4] found a high agreement between genotyping methods as long as sequencing depth was sufficiently high. This will also likely be the case when applying ACACIA instead of AmpliSAS to such datasets.

An important benefit of the Oligotyping tool in ACACIA is that unlike other clustering methods which use the entire sequence, it only uses the base pairs with the most discriminant information based on entropy analyses [13]. In the context of MHC genotyping in particular, such a strategy makes much intuitive sense, since most functional differences between MHC alleles will be within specific regions of the sequences which will contain the antigen-binding sites that are highly polymorphic as a result of strong positive selection.

### The challenge of designing primers for non-model organisms

A common approach for primer design in complex genomic regions of non-model organisms includes aligning multiple sequences of phylogenetically related species. By building primers on consensus sequences, researchers assume that oligos will amplify the target region also in the species of interest. However, knowledge about related species is often limited to very few individuals. This means that primers can be designed in regions that are polymorphic in the target species. As a consequence, certain allelic variants are not amplified and homozygosity is overestimated. Indeed, this proved to be the case in our naive primers dataset, whereby two mismatches (1st bp and 16th bp) within the forward primer (19 bp long) were sufficient to prevent the amplification of three alleles (out of 13). Interestingly, a single base pair mismatch between the second base pair of the reverse primer and the BLB1*04 allele did not prevent the amplification of this allele, although it did suffer severely from low amplification efficiency when in competition with other alleles (Figure 4). However, high sequencing depth for the naive primer dataset prevented this allele from dropping out, regardless of the genotyping pipeline used. Our study therefore highlights the importance of designing multiple primers when wishing to genotype a novel target region in non-model organisms to limit allele dropout due to primer mismatch.

### Relationship between number of alleles amplified and artefacts

By knowing the exact alleles to expect for the chicken genotypes, we were able to quantify chimeric artefacts precisely (Figure 1). There was a higher proportion of chimeric and non-chimeric artefacts in the optimal primer dataset than in the naive primer dataset. The most likely explanation for the latter is the shorter sequence for the naive primer dataset (151 bp) compared to the optimal primer dataset (241 bp). A shorter fragment reduces the number of base pairs that can be erroneously substituted and the number of breaking points for chimera formation. In addition, it is likely that the probability of incomplete elongation is inversely related to fragment length. Thus, fragment length appears to be the dominant factor predicting the proportion of artefactual reads.

As expected, the proportion of reads that were non-chimeric artefacts increased linearly as CNV increased, which can be explained simply by the fact that there is an increasing number of possible artefacts that can be generated as the number of initial template variants increases. Thus, reads that failed to be completely elongated within the PCR cycles are more likely to be erroneously elongated during the final extension step.

A more unexpected result was that the proportions of chimeras did not increase with increasing CNV, when amplifying more than 4-6 alleles. Similarly, when amplifying more than 10 alleles, the number of chimeric variants no longer increased with increasing CNV. Such saturation in chimera generation beyond a CNV threshold is likely to be a by-product of allele PCR competition. Indeed, as demonstrated by our own data (Figure 4), there is amplification bias whereby some gene variants are amplified preferentially relative to others [33, 53]. Therefore, a few gene variants (~3-6 gene variants) are preferentially amplified and most chimeras originate from these dominantly amplified variants and few chimeras are generated from the poorly amplified variants. Indeed, we found that the number of parental variants generating chimeras in our dataset did not increase with increasing CNV when amplifying more than 4-6 alleles. The non-linear relationship between chimera generation and CNV have important implications when considering sequencing depth needed to accurately genotype complex multigene system, since it suggests that linearly increasing sequencing depth for increasing CNV is not necessarily the optimal strategy. The challenges of dealing with chimeras in genotyping pipelines is discussed below in detail.

### Chimeras in genotyping pipelines

The formation of artificial chimeras during amplification is an important source of artefacts in amplicon sequencing projects [29, 52], including those with newer sequencing technologies [28]. Chimeras are challenging to identify as artefacts because they resemble real alleles generated by recombination, particularly in multigene systems under high rates of interlocus genetic exchange (“concerted evolution”), which is common in many MHC systems [6, 7, 12, 15, 18, 60]. Our results suggest that chimeras are more prevalent, harder to identify and potentially more reproducible across technical replicates than previously assumed. We expect the same to be true for similar projects with conserved, yet variable amplification targets such as the MHC.

For the optimal primer dataset, regardless of the genotyping pipeline used, allele dropout occurred in genotypes with high CNV (for ACACIA 8 out of 9 and for AmpliSAS 12 out of 14 haplotypes had a CNV < 10). For all instances bar one, allele dropout were alleles from the B21 haplotype which amplified poorly when CNV was greater than 6 (Figure 2f). Higher sequencing depth will reduce or even remove such allele dropout instances [4]. Indeed for the naive primer dataset, sequencing depth was twice as high, and there were no instances of allele dropout due to the ACACIA pipeline (all allele dropouts were due to primer mismatch). One allele erroneously called as a real variant (i.e. a false positive) by the AmpliSAS pipeline in the optimal primer dataset was actually a chimera between the BLB1*21 and BLB2*21 alleles. Furthermore, when using the AmpliSAS pipeline, 15 allele dropouts in the naive primer dataset were due to erroneous assignment of real allelic variants as chimera artefacts. Indeed, the BLB2*12 or *19 minor allele was identical to potential chimeric artefact sequences between BLB1*14 (85 possible breakpoints) and any of the following alleles: BLB2*04, BLB1*15, BLB1*19, BLB1*21 or BLB2*21 (Figure 5a.). In addition, BLB1*14 dropped out because it is identical to a chimera formed between the BLB2*02 minor and BLB2*12 or *19 alleles (33 breakpoints; Figure 5b.).

**Figure 5.**
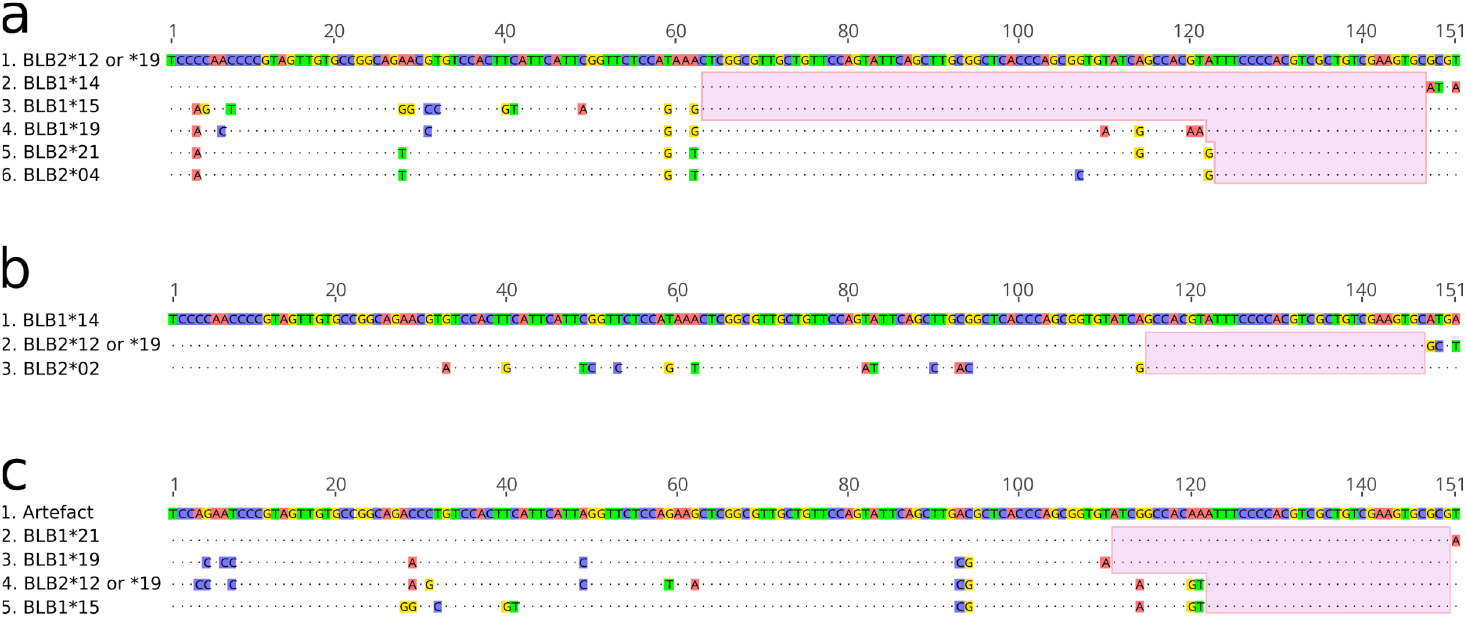
Three alignments with examples of sequences which can be classified as chimeras. The points denote identity to the first sequence in each alignment, while the differences to it are highlighted. The shaded areas indicate possible chimera-yielding breakpoints. (a) The allele BLB2*12 or *19 could be a chimera of BLB1*14 with any of the four other allele sequences depicted, in a case of multiple potential parent pairs. (b) BLB1*14 can be interpreted as a chimera between BLB2*12 or *19 minor and BLB2*02. (c) Actual chimera with multiple potential parents and a peripheral breakpoint, and therefore very similar to one of its parents.

We have identified two factors which seemed to enhance chimera formation and challenge the distinction between artefact and real allelic variants. First, the combination of multiple real “parent” sequences can yield the same chimeras, as illustrated in our examples in Figure 5a. and Figure 5b., whereby any breakpoint in the shaded areas leads to the same chimeras. Second, peripheral breakpoints (Figure 5c.) can generate chimeras that differ to parental sequences by as little as a single base pair. For instance, a chimera could be a product of the allele BLB1*21 combined with any of the other alleles shown in the alignment, with a breakpoint within the shaded area (Figure 5c.). Since the potential breaking points are at the very end of the sequence, the chimera is very similar to one of its parents (in this example, it is different from BLB1*21 by only one base). In an attempt to deal with this issue as much as possible, we changed the default settings of VSEARCH so that chimeras can be detected even if they differ from one parent by one single base. Both the “multiple parents” and the “peripheral breakpoints” issues are likely to contribute to making chimeras reproducible across replicates.

### Conclusion

Genotyping accuracy and artefacts are intrinsically linked. We have demonstrated that the ACACIA genotyping pipeline provides high allele calling accuracy and repeatability. Regardless of the pipeline used, however, users should critically assess the optimal parameters to be used. We are convinced that universal default settings for optimal genotyping accuracy cannot be achieved, since optimal parameters will depend on dataset-specific generation of artefacts. The latter, in turn, varies according to species-specific CNV, DNA quality, and the conditions of PCR (e.g. extension time, number of cycles and the polymerase used) and sequencing (e.g. quality and depth). High sequencing depth allows detecting alleles that amplify poorly in complex (multigene) systems. Furthermore simple steps prior to sequencing can greatly reduce the number of artefacts generated and improve genotyping accuracy: designing more than one PCR primer pair, reducing the number of PCR cycles, increasing PCR in-cycle extension time, and omitting the final extension step. Reducing chimera formation during PCRs is particularly critical, because they are difficult to distinguish from real alleles generated by inter-locus recombination.

## Data accessibility

Raw sequences of all datasets, example input files, suggested settings and the source code at the time of this publication are available at FigShare (https://figshare.com/projects/ACACIA/66485 and doi.org/10.6084/m9.figshare.9952520). ACACIA is freely available on the GitLab at https://gitlab.com/psc_santos/ACACIA (this paper’s code is available as a snapshot tagged as V1.0, https://gitlab.com/psc_santos/ACACIA/-/tags/V1.0), under an MIT license.

## Author contributions

MG and PS conceived the study. PS wrote ACACIA. MG did the data analysis in R. MG, PS and KM ran the allele calling workflows. KM did the AmpliSAS analysis. KW participated in and supervised the lab work. KG did the lab work. SS instigated the study and heads the lab where the work was carried out. MG and PS wrote the first draft of the paper and all authors contributed to the writing in subsequent versions.

## Supplementary material

**Supplementary Table S1.**
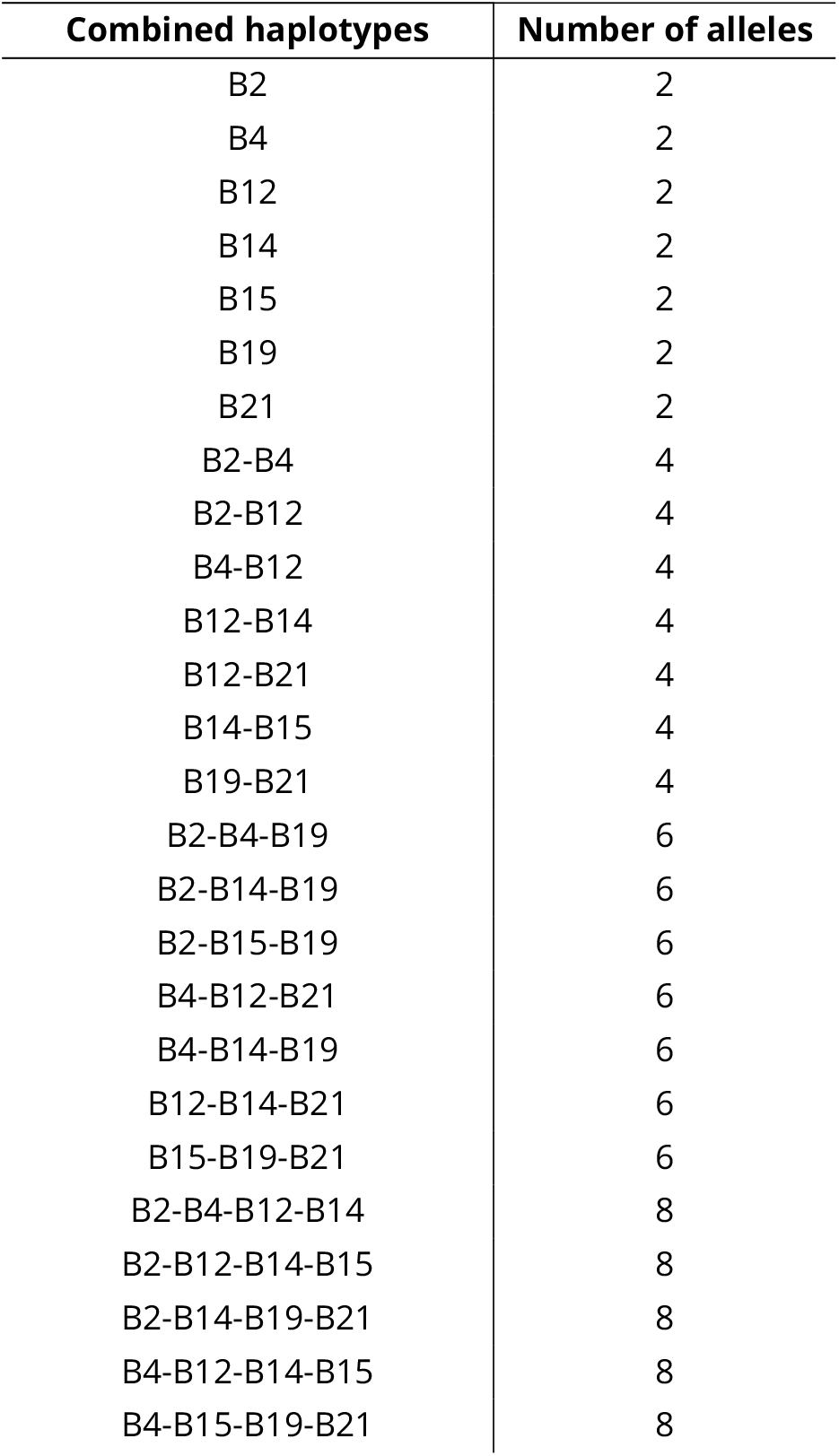
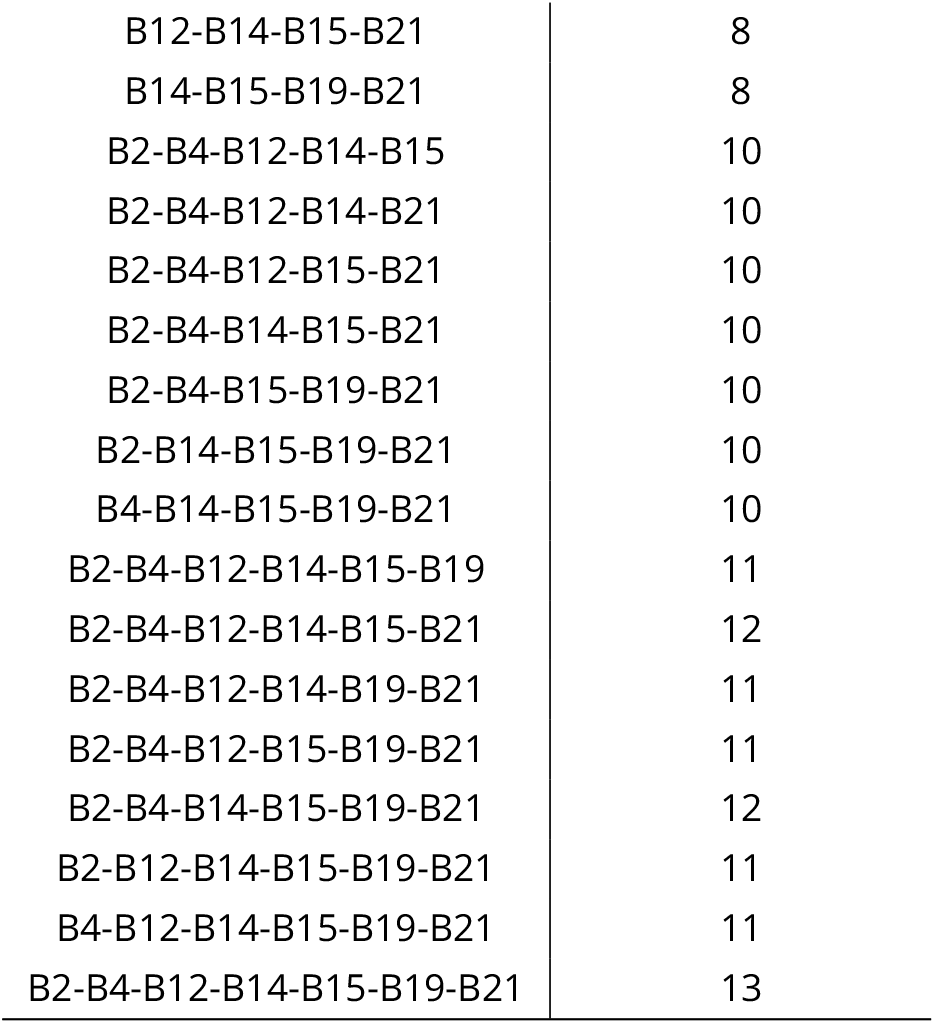
The chicken MHC B complex haplotypes and combined haplotypes which formed experimental genotypes with varying copy number variation (CNV).

| Combined haplotypes | Number of alleles |
| --- | --- |
| B2 | 2 |
| B4 | 2 |
| B12 | 2 |
| B14 | 2 |
| B15 | 2 |
| B19 | 2 |
| B21 | 2 |
| B2-B4 | 4 |
| B2-B12 | 4 |
| B4-B12 | 4 |
| B12-B14 | 4 |
| B12-B21 | 4 |
| B14-B15 | 4 |
| B19-B21 | 4 |
| B2-B4-B19 | 6 |
| B2-B14-B19 | 6 |
| B2-B15-B19 | 6 |
| B4-B12-B21 | 6 |
| B4-B14-B19 | 6 |
| B12-B14-B21 | 6 |
| B15-B19-B21 | 6 |
| B2-B4-B12-B14 | 8 |
| B2-B12-B14-B15 | 8 |
| B2-B14-B19-B21 | 8 |
| B4-B12-B14-B15 | 8 |
| B4-B15-B19-B21 | 8 |
| B12-B14-B15-B21 | 8 |
| B14-B15-B19-B21 | 8 |
| B2-B4-B12-B14-B15 | 10 |
| B2-B4-B12-B14-B21 | 10 |
| B2-B4-B12-B15-B21 | 10 |
| B2-B4-B14-B15-B21 | 10 |
| B2-B4-B15-B19-B21 | 10 |
| B2-B14-B15-B19-B21 | 10 |
| B4-B14-B15-B19-B21 | 10 |
| B2-B4-B12-B14-B15-B19 | 11 |
| B2-B4-B12-B14-B15-B21 | 12 |
| B2-B4-B12-B14-B19-B21 | 11 |
| B2-B4-B12-B15-B19-B21 | 11 |
| B2-B4-B14-B15-B19-B21 | 12 |
| B2-B12-B14-B15-B19-B21 | 11 |
| B4-B12-B14-B15-B19-B21 | 11 |
| B2-B4-B12-B14-B15-B19-B21 | 13 |

## Acknowledgements

MG was supported by a DFG grant (DFG Gi 1065/2-1). We are very grateful to Jim Kaufman and his lab members for providing the chicken DNA samples used in this study and for his comments on a previous version of this work. Version 3 of this preprint has been peer-reviewed and recommended by Peer Community In Evolutionary Biology (https://doi.org/10.24072/pci.evolbiol.100092). We thank the reviewers Thomas Bigot, Helena Westerdahl and Sebastian Ernesto Ramos-Onsins and the recommender François Rousset for their comments which improved our manuscript.

## Conflict of interest disclosure

The authors of this preprint declare that they have no financial conflict of interest with the content of this article. François Rousset is the recommender for PCI Evolutionary Biology.

## Notes

#### Summary of Updates

This article has been peer-reviewed and recommended by Peer Community In Evolutionary Biology

https://gitlab.com/psc_santos/ACACIA

https://figshare.com/projects/ACACIA/66485

